# Microplastic ingestion by a herring *Opisthonema sp.*, in the Pacific coast of Costa Rica

**DOI:** 10.1101/670679

**Authors:** Luis Bermúdez-Guzmán, Crista Alpízar-Villalobos, Johan Gatgens-García, Gabriel Jiménez-Huezo, Marco Rodríguez-Arias, Helena Molina-Ureña, Javier Villalobos, Sergio A. Paniagua, José R. Vega-Baudrit, Keilor Rojas-Jimenez

**Affiliations:** Escuela de Biología, Universidad de Costa Rica, 11501-2060, San Pedro, San José, Costa Rica; Centro de investigación en Ciencias del Mar y Limnología (CIMAR) Universidad de Costa Rica, San Pedro, San José, Costa Rica; Laboratorio Nacional de Nanotecnología LANOTEC-CeNAT-CONARE, 1174-1200, Pavas, San José, Costa Rica

**Keywords:** Plastic pollution, *Opisthonema*, marine pollution, Tropical Eastern Pacific

## Abstract

Despite there is a growing interest in studying the presence and effects of microplastics (MP) in fishes and other aquatic species, knowledge is still limited in tropical areas. In this study, we examined the presence of MP in the gastrointestinal content of 30 filter feeders of thread herring, *Opisthonema* complex (Clupeiformes: Clupeidae) from the Central Pacific coast of Costa Rica. We detected the presence of MP in 100% of the individuals with an average of 36.7 pieces per fish, of which 79.5% were fibers and 20.5% particles. To our knowledge, this is the first study in Costa Rica that demonstrates the presence of MP in planktivorous fishes. The effects of microplastics ingestion by *O. libertate* and its transit through aquatic food webs should be studied in greater detail, with greater number of sampling points at different times of the year. However, our work confirms that contamination by microplastics is having direct effects on the marine life of Costa Rica.

**Capsule:** This is the first multidisciplinary study in Costa Rica demonstrating the presence and nature of microplastics in the digestive tract of planktivorous fish.

## Introduction

Pollution by plastic in the oceans was reported since the second half of the 20^th^ century (Carpenter and Smith, 1972; Shiber, 1979), but the scientific investigation of its implications in marine life has been addressed until recently (Law, 2017). In Costa Rica, few efforts have been made to determine the presence of microplastics at different marine trophic levels, even though it is estimated that more than 500 metric tons of solid waste is discarded per day.

Microplastics are defined as particles with a size less than 5 mm that can enter into the environment by a primary form (cosmetic fragments and clothing fibers) or secondary, generated from the decomposition of larger plastic objects by means of ultraviolet (UV) photo-degradation, wave action and physical abrasion (Martin et al., 2017). The chemical composition of these particles depends on different monomers that are used for their production, among which polyethylene, polypropylene, and polystyrene are the most abundant (Güven et al., 2017).

As the size of these particles overlaps with the size of zooplankton organisms, planktivorous fish can ingest microplastics directly (Law and Thompson, 2014) or through feeding zooplankton that has previously ingested microplastics (indirectly) (Cole et al., 2011). In this line, (Setälä et al., 2014) evidenced the ingestion of 10 μm polystyrene microspheres in all the planktonic organisms studied, including shrimp, copepods, cladocerans, rotifers, and polychaete larvae It also demonstrated the transfer of plastic microparticles through planktonic guilds from lower trophic levels (mesozooplankton) to a higher level (macrozooplankton).

Among the organisms that could potentially ingest microplastics are filter feeders of the Clupeidae family, whose feeding depends on planktonic life stages (Lozano, 1979). Recent studies have found the presence of microplastics in the digestive tract of clupeid fish (Ory et al., 2018; Tanaka and Takada, 2016). These fish represent a good study model since their short life, and their tendency to group in homogeneous schools provide an updated image of the amount and type of microplastic present in the marine landscape (at any given moment).

Due to the enormous increase registered over the last century, both in the production of plastics and in their presence in the oceans worldwide (Ivar do Sul and Costa, 2014), it is necessary to know their scope in marine ecosystems in order to determine its impact and take mitigation measures of possible damages. The objective of this investigation was to determine the incidence of microplastics in the gastrointestinal content of *Opisthonema* sp., a planktivorous fish in the Pacific Coast of Costa Rica and a model for biomonitoring microplastic pollution in the marine ecosystem.

## Materials and Methods

### Study site

Samples from commercial purse-seine catches, collected by the Costa Rican semi-industrial sardine fleet, were obtained in October 2018 in the province of Puntarenas, Central Pacific of Costa Rica (09’49.’242N, 084’42.087 W) (**Fig. 1**). Three species of the *Opisthonema* complex (*O. libertate, O. bulleri, O. medirastre*) sustain the sardine fishery of Costa Rica, although the Pacific thread herring, *O. libertate* tends to be most abundant in their mixed schools during the sampling period (Vega-Corrales, 2010). Non-eviscerated whole specimens were transported fresh from the landing dock to the Center for Research in Marine Sciences and Limnology (CIMAR) of the University of Costa Rica (UCR).

**Figure 1.**
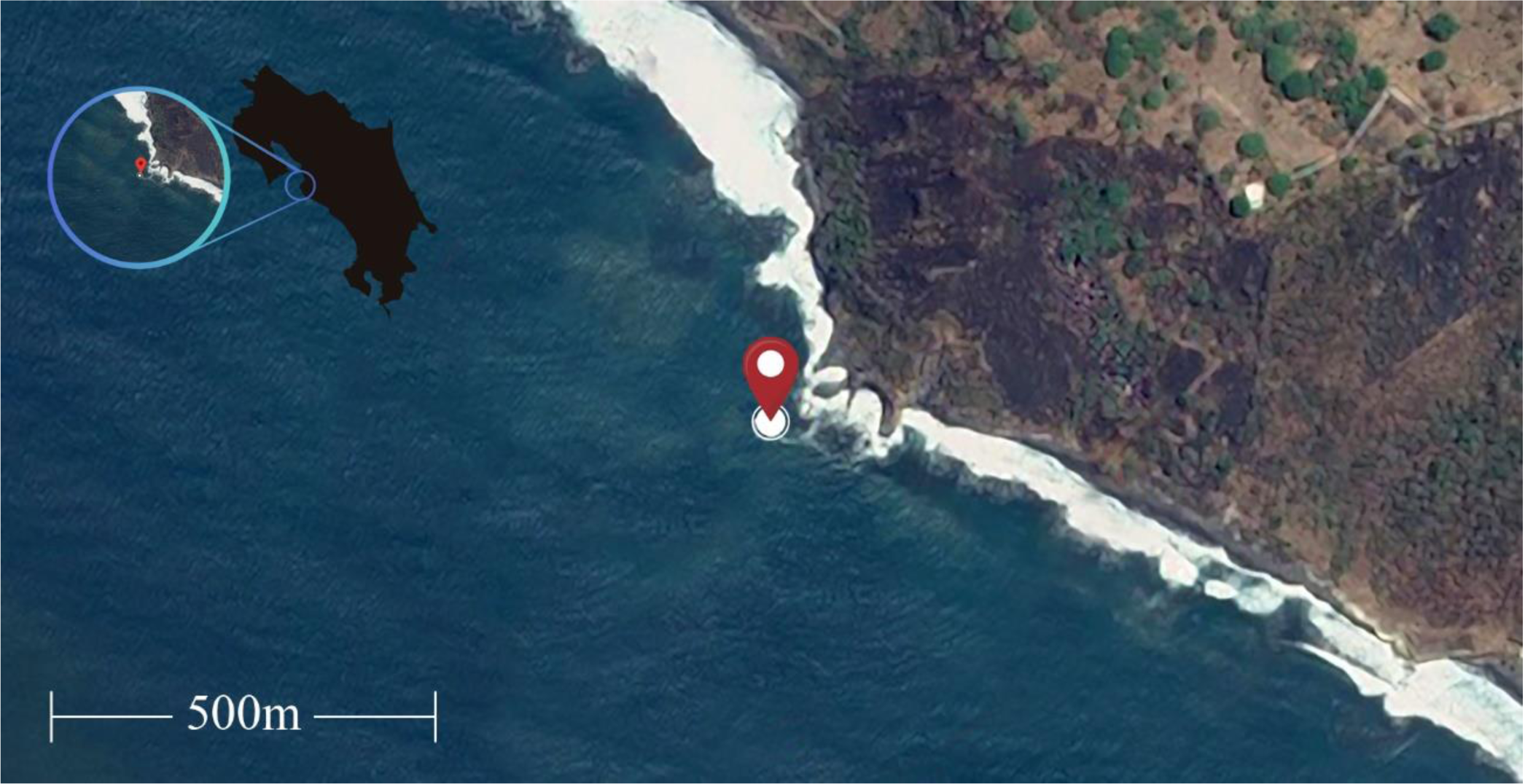
The geographic location of the sampling site (in red) in the central Pacific coast of Costa Rica.

### Analysis of samples

Thirty individuals of *Opisthonema* complex (*O. libertate, O. bulleri, O. medirastre*) underwent a series of biometric analyses in CIMAR laboratories. We measured the standard length (SL), fork length (FL) and total length (TL) of each fish. Subsequently, their total and eviscerated weights were determined on a Mettler PJ360 DeltaRange® granatary scale. Then, each fish was opened with dissection scissors tracing a straight line in the belly from the anus to the preopercular area, extracting the organs associated with the gastrointestinal tract. The gastrointestinal tract was isolated from the mesentery and other structures in order to measure and weigh it. Each specimen was sexed by examining the gonads.

### Processing of the intestinal tract

A longitudinal section was made through the whole tract to obtain the gastric content, then it was deposited in a filter paper. The tracts were washed with filtered distilled water and filtered to obtain all the organic matter and the microplastics. The material was dissolved in KOH 10% (previously pre-filtered though 0.2 µm) for the degradation of the organic matter. This mixture was left standing for a minimum period of 48 hr in separate glass bottles.

After the incubation, each sample was vacuum-filtered using a Watman celulose filter paper. The filters were dried under a 100 Watts incandescent lamp and subsequently analyzed under Motic® brand DM-143 stereoscopes. The material obtained were separated into fibers and particles. As a negative control, a portion of the KOH was incubated and later filtered to analyze it under the stereoscope and determine if any MP comes from our protocol.

### Fiber and particle analysis

Scanning electron microscopy (SEM): The particles isolated from the gastrointestinal tract of *O. libertate* were mounted on carbon tape and sputtered with gold using a Denton Vacuum Desk V sputter system at 20 mA for 300 s. Images were taken using a JSM-6390LV (JEOL, Tokyo, Japan) SEM, with an accelerating voltage of 20 kV, under high vacuum. Energy-dispersive X-rays (EDX)were measured with liquid nitrogen cooled Inca X-sight Si detector (Oxford Instruments). EDX data was analyzed with Inca Suite version 4.08.

Fourier-transform infrared spectroscopy with attenuated total reflection (FTIR-ATR): The spectra were collected in the range 4000-500 cm^−1^, using a Nicolet 6700 Thermo Scientific spectrophotometer with a diamond ATR crystal.

Differential scanning calorimetry (DSC). The analysis of *ca.* 1 mg of particles was performed in a Q200 TA Instruments calorimeter, under nitrogen atmosphere in the temperature range - 80 – 200 °C, at a rate of 20 °C min^−1^.

### Statistical analysis

Data processing and statistical analysis were performed in R (R Core Team, 2018). Visualizations were generated with the program ggplot (Wang et al., 2017) (Wickham, 2016). The statistical differences in the number of microplastic particles were estimated using the non-parametric Kruskal-Wallis test.

## Results

The most notorious result of this work is that microplastics were found in 100% of the individuals. We counted a total of 1100 pieces in the 30 individuals analyzed, of which 20.5% were classified as particles and 79.5% were fibers. The average number of pieces per fish was 36 (range 32 to 42) where the average number of fibers was 29 (range 25 to 34), and the average number of particles was 5 (range 6 to 10). These results represent the first stage of our work; therefore, sample number should be increased in subsequent studies.

From the 30 specimens of *Opistonema* sp., five were females and 25 males. The measurements of the main characteristics of the fishes (including standard, fork, and total length; length of the gastrointestinal tract; average weight with and without evisceration; average weight of the full and empty tract) grouped by sex are shown in **Fig. 2**. We also observed a trend of higher total number of MPs in females than in males, but results were not statistically significant (**Fig. 3**). Additional studies with more samples from each sex are required to determine if females have higher ingestion rates of microplastics than males. We did not find any other apparent association between the variables measured and the ingestion of microplastics. This could be relevant since it shows that this species can be used as a standard model for the biomonitoring of microplastic in the sea.

**Figure 2.**
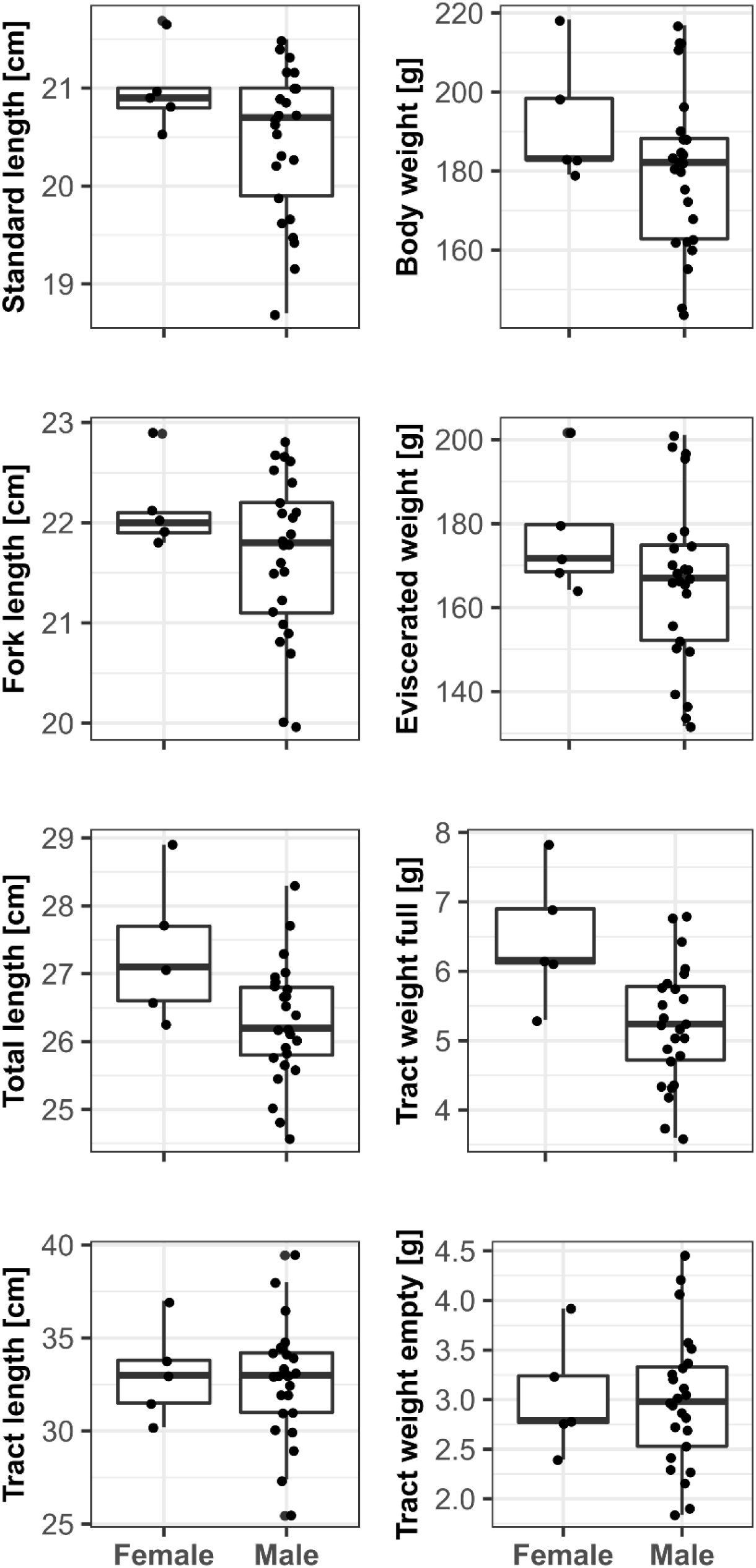
Measurements of the main characteristics of the samples of *O. libertate* grouped by sex.

**Figure 3.**
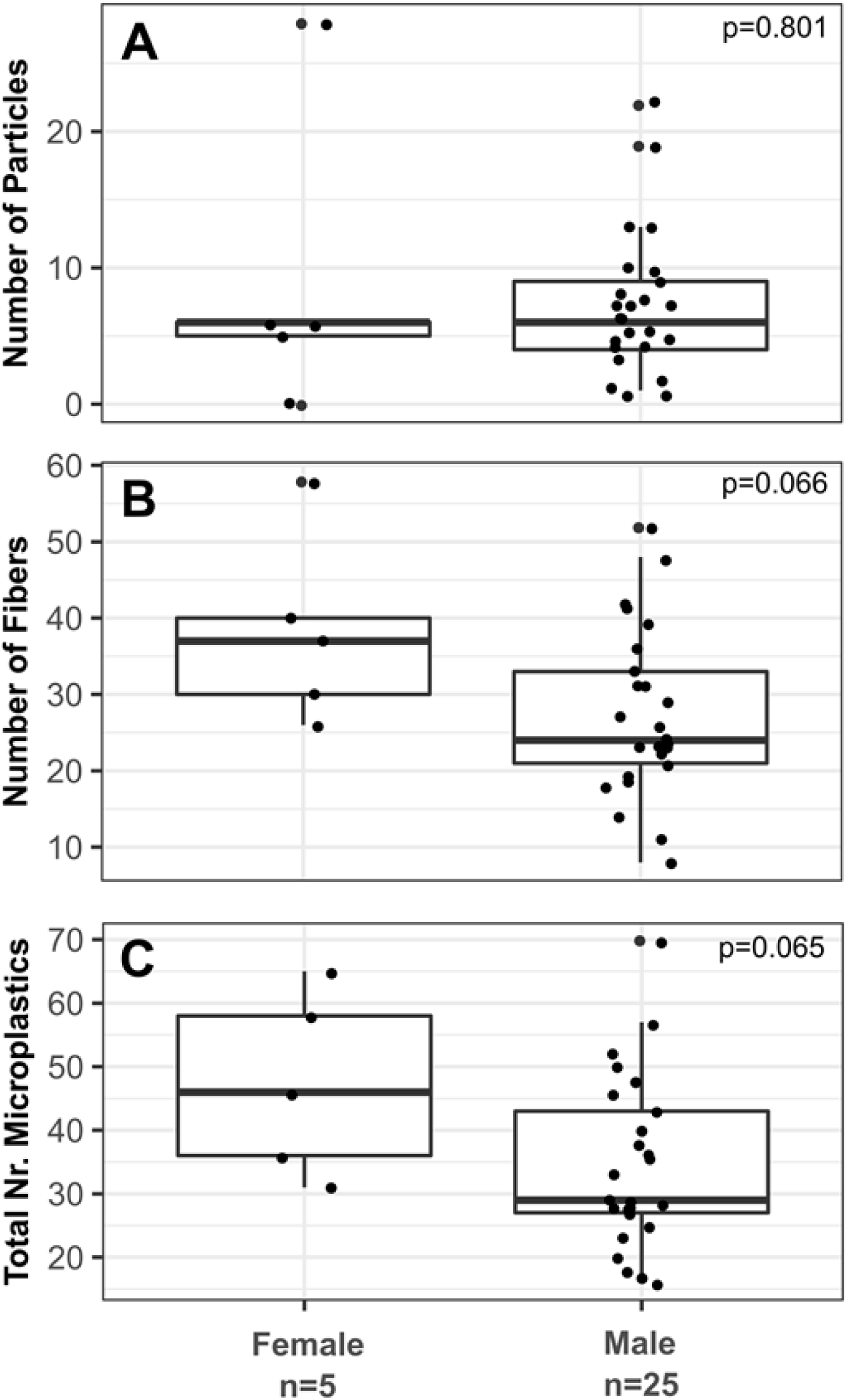
Number of particles of microplastics identified in the gastrointestinal tract of *O. libertate* and grouped by sex.

The main shapes and sizes of the particles and fibers found in the digestive tract of the fish were photographed using a stereoscope **(Fig. 4)**. In addition, SEM was used to determine which of the structures found corresponded to MP. The analysis showed three types of structures to be screened (**Fig. 5**): fibers (tagged as A), incrustations (attached to fibers, tagged as B) and a couple of standalone particles (tagged as C). EDX spectroscopy was run on selected structures to determine the chemical composition. When no incrustations were observed around the fibers, only C and O were detected, which agrees with organic substances. Incrustations typically added Ca, K and Si to the composition. Similar crusts have been reported on other MP studies (Wang et al., 2017). A partially mineralized microorganism is tagged with an asterisk symbol. Other SEM images showing mineraloid incrustations in the fibers are available in supplementary information. The standalone particles tagged as (C) in **Fig. 5** were inorganic, with no Carbon detected (see EDX results for rod-like structure as an example-the other standalone structure tagged as (C) was an iron-rich aluminosilicate). This is relevant since, in many cases, the classification of microplastics in fibers and particles by optical microscopy could generate an overestimation of MP if their chemical nature is not verified by other methods.

**Figure 4.**
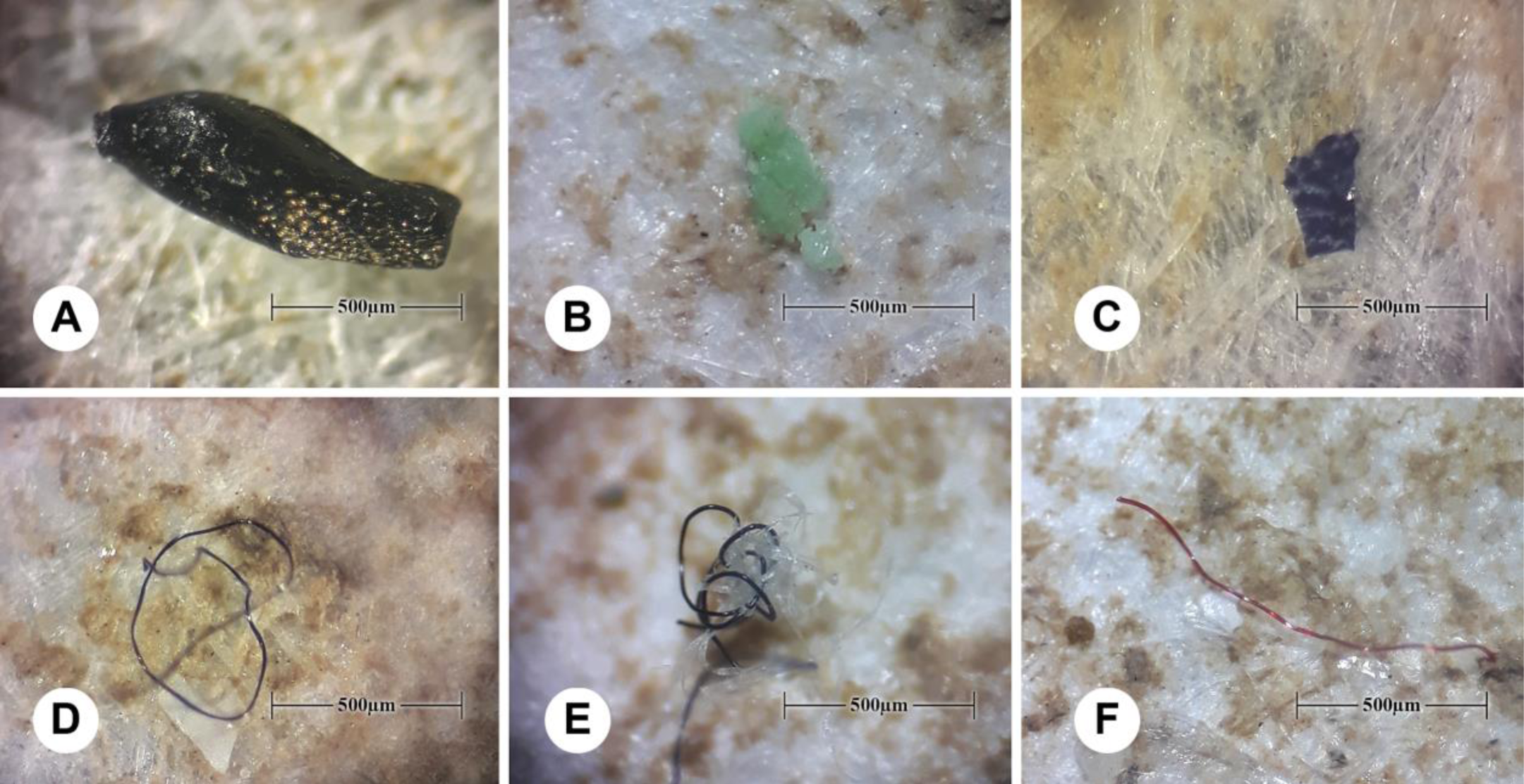
Micrographs of microplastics found in the digestive tract of *O. libertate*. The images A, B, and C correspond to particles, and the images D, E, and F correspond to fibers.

**Figure 5.**
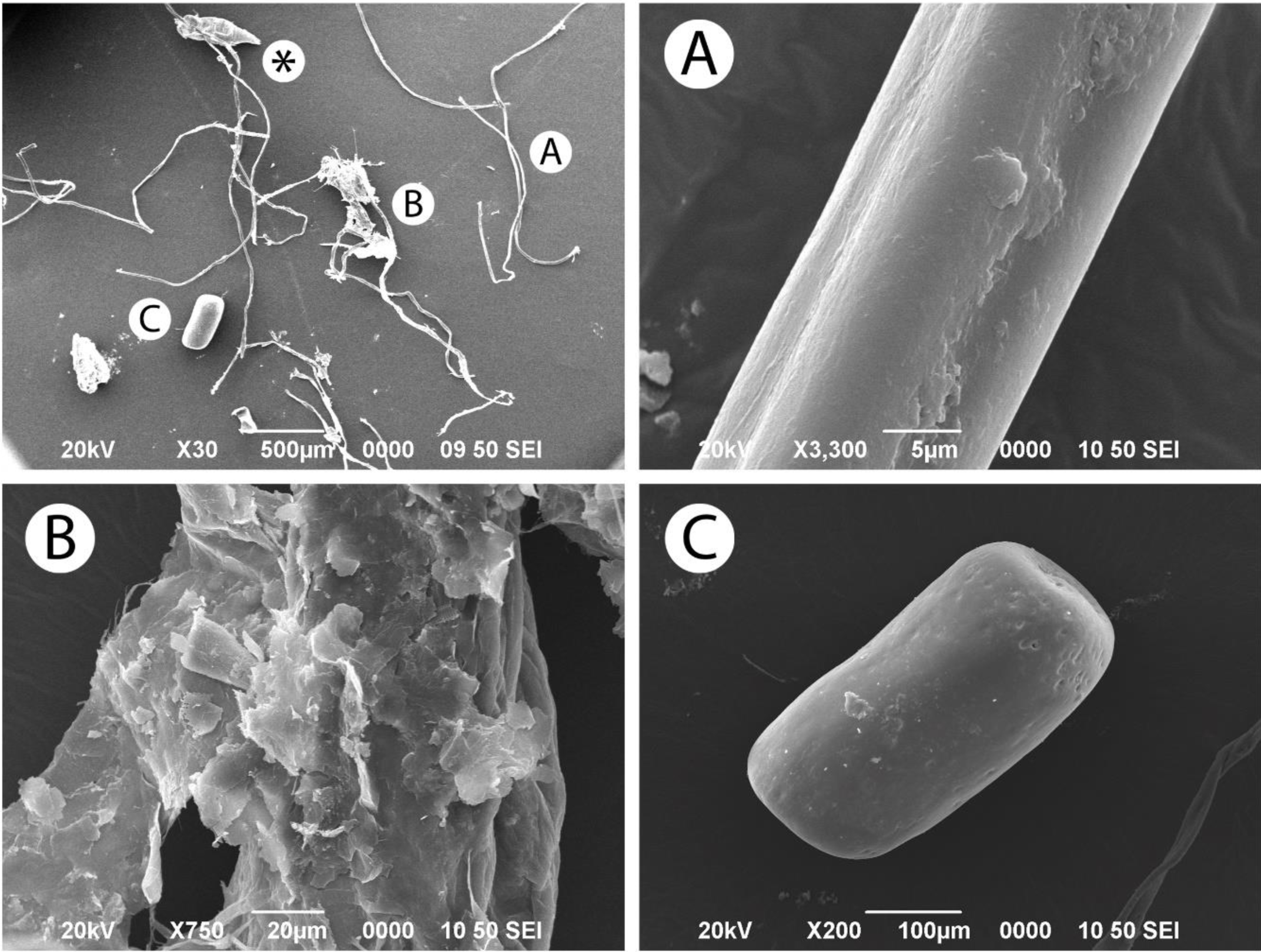
SEM images of contents in the digestive tract of *O. libertate* and their % weight of each element by Energy Dispersive X-ray Spectroscopy (EDX). The low magnification view on the left shows different types of solids that can be categorized as (A) fibers, (B) incrustations, and (C) standalone particles. Fiber in A showed a composition by weight % of Carbon (60±2) and Oxygen (40±2). In B, EDX showed a variable composition; Aluminum (4±1), Calcium (10±2), Carbon (35±11), Potassium (4±1), Oxygen (38±7) and Silicon (9±2). Particle in C, showed the following composition: Calcium (57±4), Magnesium (5±1) and Oxygen (38±5).

FTIR-ATR and DSC were used to determine the types of polymers contained in the samples. The FTIR-ATR spectrum (**Fig. 6)** shows typical signals for thermoplastic polyolefins, such as polyethylene or polypropylene: the CH_2_ asymmetric stretching (2917 cm^−1^) and CH_2_ symmetric stretching (2839 cm^−1^), the CH_3_ symmetric deformation (1376 cm^−1^) and the CH_2_ bending deformation (1456 cm^−1^) (Gulmine et al., 2002). Moreover, there are some weak signals around 3300 cm^−1^ and 1700 cm^−1^, suggesting a small grade of oxidation of the polymer.

**Figure 6.**
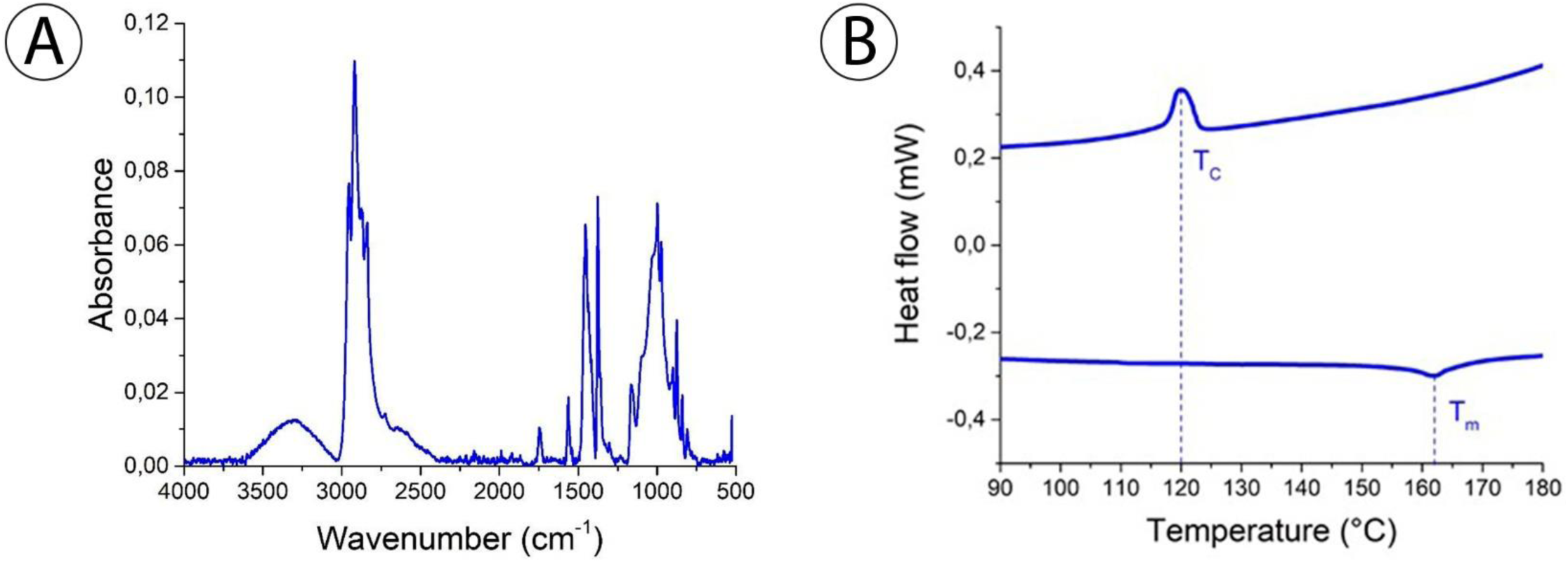
**A)**-ATR spectrum for contents in the digestive tract of *O. libertate*. Range:500-4000 cm^−1^. **B)** DSC graph: heat flow (mW) versus temperature (°C) for *ca* 1 mg of contents in the digestive tract of *O. libertate* under nitrogen atmosphere. Heating/cooling rate: 20 °C min^−1^.

The DSC curve (**Figure 6.B**) shows two main signals, an exothermic signal at 120 °C and an endothermal signal at 162 °C, which represent the crystallization temperature (Tc) and the melting point (TM) of the sample, respectively. As many thermoplastics, polypropylene has a wide range of melting and crystallization temperatures, depending on molecular weight and presence of functional groups. (Majewsky et al., 2016) reported the endothermic peaks (TM) by DSC for typical polymers in order to identify them in MP samples. The reported TM for polyethylene was 101 °C, 164 °C for polypropylene, and 250 °C for polyethylene terephthalate (PET).

The FTIR-ATR and DSC results of the batch analyzed suggest the presence of polypropylene in our MP samples, which is expected since it is widely used in packaging, labeling, containers, and others. It is important to mention that these results do not exclude the possible presence of other less concentrated polymers in the sample. In addition, the proportion of the sample analyzed was very low compared to the total number of pieces that were found.

## Discussion

### The utility of Opistonema as a model for biomonitoring

To our knowledge, this is the first study in Costa Rica demonstrating the presence of microplastics in the digestive tract of planktivorous fish of the Clupeidae family. Although there are no other published studies in the country to compare with, this is the first study in the region showing 100% of the samples containing microplastics in their intestinal tracts and also accounting for the highest number of microplastic pieces per fish (Espinoza and Bertrand, 2008; Ory et al., 2018, 2017; Tanaka and Takada, 2016). Despite the low number of samples analyzed, this study sets a precedent and calls for the continuous monitoring of the effects of microplastic contamination on the coasts of the region.

The three measures of length showed little variation among the fish, and we found no correlation between average weight and length with the number of microplastics. The homogeneous biometric characteristics of the mixed schools of *Opisthonema* allows us to suggest that this species complex could be used as a model biomonitoring studies of MP pollution. In addition, the feeding type of this species makes it more prone to the intake of microplastic, since, by suction, they cannot discriminate the presence of microplastic (Moore et al., 2002, 2001). This type of feeding could explain the high number of pieces found per fish as well as the small differences between individuals. In this regard, (McNeish et al., 2018) concluded that filter fish tend to have more particles and plastic fibers than others with a different type of feeding, due to the trophic transfer of the plastic elements consumed by the prey.

As we found fibers of different colors, it is possible that there is no discrimination for this characteristic (Tanaka and Takada, 2016). In contrast (Ory et al., 2017) reported that in *Decapterus muroadsi* (Carangidae) the microplastic capture could be due to a confusion between the color of the particle and the color of its prey. The fact of having found so many pieces in short-lived fish might serve as a base for estimating the number of pieces that are floating in the photic zone, which is where this species usually inhabit. To have a complete picture, it will be necessary to analyze water samples from the nearby areas as well (Güven et al., 2017).

### Comparison between the Pacific coast of different countries

Different studies performed on the Pacific coast of other countries have shown the presence of MP in the different levels of the trophic chain (Law and Thompson, 2014). For example, research conducted in the Pacific coast has shown the existence of MP in filter fish from Japan (77%), Chile (Easter Island) (80%) and in California, United States (35%). However, in countries such as Peru, Colombia, and Panama, no microplastics were detected ((Boerger et al., 2010; Espinoza and Bertrand, 2008; Ory et al., 2018, 2017; Tanaka and Takada, 2016) (**Table I**). Since *O. libertate* was also used in Colombia for biomonitoring MP contamination, we propose this species as a model for comparison between countries in the Pacific coast of the region. An explanation for the differences between countries in the region can be related to the effects of the marine currents, that transport the microplastics, and that convergence near to Central America and North America (Law, 2017). However, we consider that the proximity to urban areas with a high degree of pollution is the most important agent.

**Table 1.** Comparison of the percentage of fish that had microplastic and their respective average in their digestive tract, according to the country (year) and species of study.

| Country | Specie | Number of fishes | Percentage of fish with MP (%) | Average of MP per fish (±s.d.) | Total of MP |
| --- | --- | --- | --- | --- | --- |
| Costa Rica (2018) | *Opisthonema libertate* | 30 | 100 | 36.7(±0.86) | 1101 |
| Easter Island (2017) | *Decapterus muroadsi* | 20 | 80 | 2.5(±0.4) | 48 |
| Japan (2015) | *Engraulis japonicas* | 64 | 77 | 2.3(± 2.5) | 150 |
| Colombia (2016) | *Opisthonema libertate* | 27 | 0 | 0(±0) | 0 |
| Panama (2018) | *Cetengraulis mysticetus* | 10 | 0 | 0(±0) | 0 |
| Peru (2018) | *Engraulis ringens* | 40 | 0 | 0(±0) | 0 |

### Implications in marine life

Floating plastics can be transported to greater depths by increasing the density induced by biofouling and can be ingested by migratory species. Many of these processes have been demonstrated in laboratory and field experiments, but their rates at a global scale remain unknown (Law, 2017). The latter could also present a risk to other organisms of different trophic levels, such as crustaceans or birds that feed on other fish. About other implications of the MP for the marine species, it has been proven that MP releases toxic substances that include residual monomers, plasticizers, coloring agents, among other additives, that can be ingested and produce bioaccumulation (Worm et al., 2017). In laboratory studies, it was found that MP particles ingested by fish of the Clupeidae family (*Alosa fallax*) passed from the digestive system to the circulatory system and later to other organs (Neves et al., 2015). Likewise, in other experiments, it has been shown that the exposure of reef fish to water sources that had previously been exposed to polypropylene bags raises the levels of nonylphenol in the fish, which led to their short and long-term death (Worm et al., 2017). Another consequence of the presence of MP in the digestive tract of fish could include choking, histological damage and alteration of the microbiome (Batel et al., 2018; Jin et al., 2018; Karami et al., 2017).

Plastic debris can also harbor pathogens that are often associated with disease outbreaks, e.g., in coral reefs, since microbial communities can colonize microplastics. An example of this is the bacteria of the genus *Vibrio* (Zettler et al., 2013), an opportunistic pathogenic bacterium known to cause coral diseases worldwide (Lamb et al., 2018).

## Conclusions

This study represents an emerging research field for Costa Rica as a response of the efforts made in the region to characterize the presence of microplastics in marine life, and its possible ecological implications. Although this is a small study, our results help to integrate existing information on MP contamination in marine life. The validation of *Opistonema* sp. as a model species in the MP biomonitoring will require more studies. It is also necessary to replicate this type of studies systematically in different points of the Pacific coast and at different times of the year, to better understand the effect of local and regional currents on the dynamics of microplastic masses. In the Caribbean region, it is also necessary to carry out this type of research to compare the state of the two Costa Rican coasts.

It is important to develop strategies that attempt to identify the possible physiological effects of MP at different levels of marine life. In the case of our model species, it would be necessary to investigate if there is any involvement at the histological level, at the metabolic level or even in the microbiome (dysbiosis) due to the high presence of MP.

## Acknowledgments

We thank the Centro de Investigación en Ciencias del Mar y Limnología (CIMAR) of the University of Costa Rica. We also thank Jeffrey Sibaja, Cindy Fernandez, Juan José Alvarado, and Gerardo Umaña for their valuable help during the development of this investigation. KRJ was partially supported by project B8-297 of the University of Costa Rica (UCR).

We thank the National Nanotechnology Laboratory (LANOTEC) of the National High Technology Center (ceNAT), as well as Ethel Sanchéz Chacón.

## Supplementary information

**Supplementary Figure 1.**
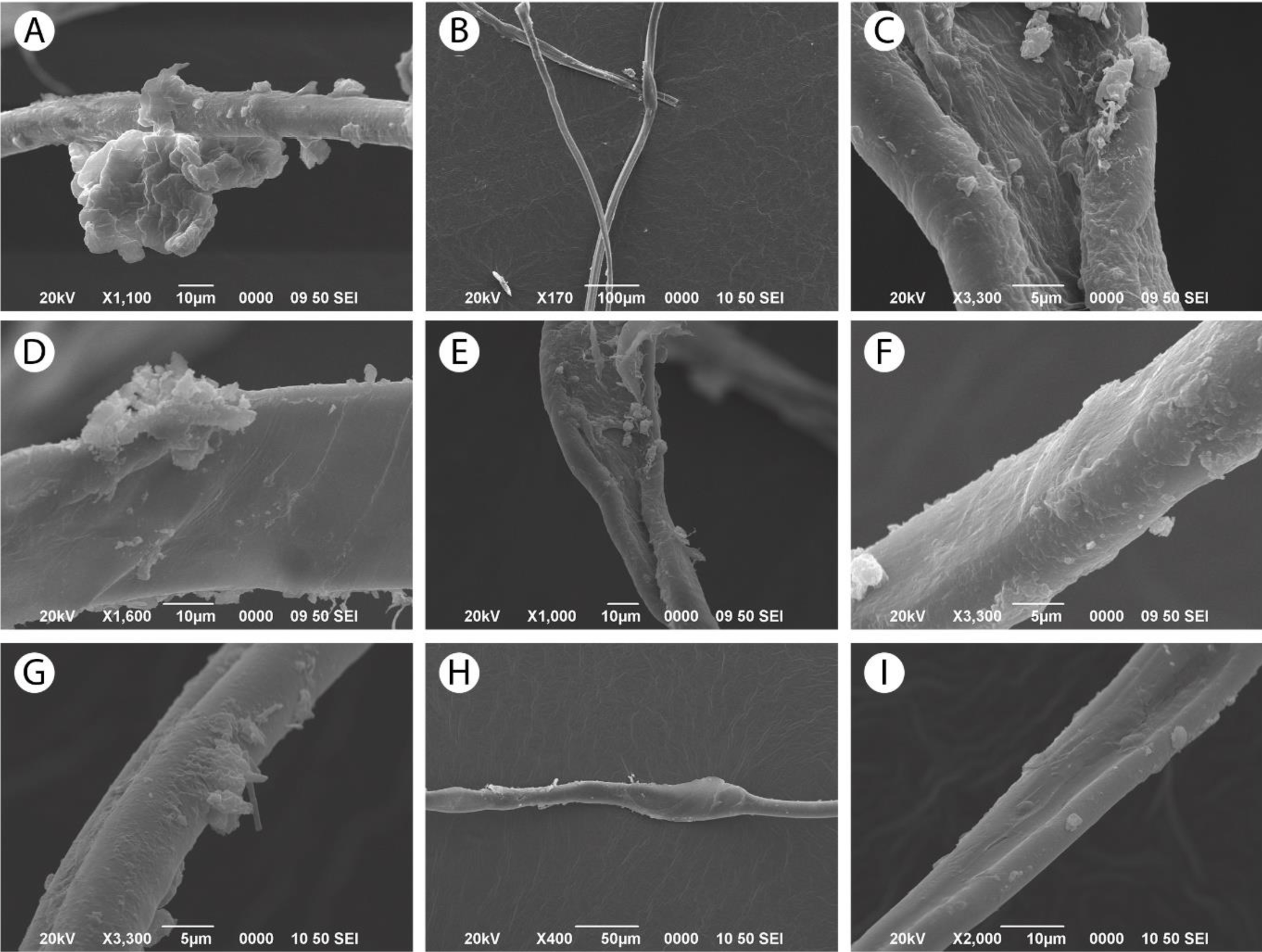
SEM images of fibers found in the digestive tract of *O. libertate.* The presence of incrustations in the fibers was evident at ∼3000X. It was possible to confirm the mineraloid nature of the incrustations based on the composition obtained by Energy Dispersive X-ray Spectroscopy (EDX). Incrustations are more evident in A, C, D, G.

